# The effects of combining focus of attention and autonomy support on shot accuracy in the penalty kick

**DOI:** 10.1101/560813

**Authors:** Hubert Makaruk, Jared M. Porter, Jerzy Sadowski, Anna Bodasińska, Janusz Zieliński, Tomasz Niźnikowski, Andrzej Mastalerz

## Abstract

The penalty kick is of great importance in the sport of soccer. Therefore, the aim of this study was to test predictions of the OPTIMAL theory and identify key attentional and motivational factors that impact the accuracy of the penalty kick. The following six groups of skilled participants performed penalty kicks following instructions that directed their focus of attention or impacted their autonomy support: external focus with autonomy support (EF/AS), external focus alone (EF), internal focus with autonomy support (IF/AS), internal focus alone (IF), autonomy support alone (AS) and control (C) groups. The analysis showed that the EF/AS group demonstrated better kicking accuracy relatively to the IF/AS, IF and C groups, but there were no significant differences between the EF/AS and EF or AS groups. Interestingly, the EF/AS group showed higher self-efficacy compared to the EF, IF/AS, IF and C groups. The finding suggest that a combination of attentional and motivational factors may produce benefits in motor performance.

## Introduction

For two decades, research findings have consistently demonstrated that adopting an external focus of attention improves motor performance compared to attempts performed when using an internal focus of attention (for a review see [1]). Studies have also reported that granting the mover autonomy (e.g., self-control, enhanced expectancy) over some aspect of practice also improves motor performance [2]. Recently, Wulf and Lewthwaite [2] proposed the Optimizing Performance Through Intrinsic Motivation and Attention for Learning (OPTIMAL) theory as an alternative way to explain how practice effects such as attention and autonomy support influence motor performance and learning. This theory proposes that practice environments which encourage the mover to utilize an external focus of attention, promote autonomy support by way of self-controlled practice methods, and link movement goals with actions increase intrinsic motivation which then facilitates motor learning.

The predictions of the OPTIMAL theory have been tested in contemporary studies. For example, Pascua et al. [3] examined undergraduate students’ throwing in four different practice conditions: external focus with enhanced expectancy, external focus only, enhanced expectancy only and a control group that did not receive any attention directing instructions or enhanced expectancy. The authors reported that the combination of an external focus and enhanced expectancy resulted in additive effects in throwing accuracy during retention and transfer tests. Similarly, findings of another study [4] showed that matching autonomy support and enhanced expectancies provided the highest accuracy score compared to all other experimental conditions when throwing a ball. Whereas using these factors alone showed intermediate scores compared to the control group, which had the lowest scores on the retention and transfer tests. Wulf et al. [5] reported that using an external focus combined with autonomy support yielded the superior accuracy compared to other practice groups that used each factor separately. In each of these studies, the pairing of two factors (e.g., external focus and autonomy support) had an enhancing effect on motor learning compared to practicing with only an external focus or with autonomy.

In the present study we were interested in testing how quickly the combination of autonomy support and focus of attention affected performance compared to only giving the mover autonomy or altering their focus of attention during the execution of a well learned skill. This is important for both theoretical and practical reasons. Building on previous research, the primary aim of this study was to investigate whether combining an external focus of attention with autonomy support produced a motor performance advantage relative to these factors being adopted alone. In addition to pursuing this aim, we wanted to examine the predictions of the OPTIMAL theory and gain a deeper understanding of how combining these factors during motor performance impacted movement accuracy. If the predictions of the OPTIMAL theory are correct, then the combination of autonomy and an external focus of attention should result in enhanced movement accuracy when performing a well learned skill relative to practice conditions that do not combine these factors. The present experimental design involved six conditions: external focus with autonomy support, external focus alone, internal focus with autonomy support, internal focus alone, autonomy support alone and a group not receiving focus directing instructions or autonomy support (i.e., control condition).

The penalty kick was chosen as a suitable task for this study for two reasons. First, because of the importance of the penalty kick in determining the outcome of soccer matches. A recent study [6] reported that soccer teams awarded penalty kicks during match play won 52%, drew 30% and lost 18% of those matches. The authors also indicated that the chances of winning a match increased to 61% if the penalty kick was scored, but decreased to 29% if the penalty shot was missed. Additionally, teams participating in either the World Cup or European Championship final match had roughly a 50% chance of being involved in a penalty shootout during the tournament. It is also worth pointing out that the conditions of performing the penalty kick are decisively more favorable for the penalty kicker rather than the goalkeeper. Meaning that the goalkeeper must defend a relatively large goal area (7.32 x 2.44 m) from a relatively short distance of 11 m. With this in mind it is interesting to note that approximately 20-40% of penalty kicks completely miss the goal [7–9]. These findings clearing indicate if a soccer player can improve his or her penalty kick shooting accuracy it has a significant impact on determining the outcome of the game, especially in tournament matches.

A second reason we utilized the penalty kick in the present study is that the accuracy of the penalty kick may be improved when suitable instructions are provided [10,11]. Researchers suggest that the skill of hitting optimal areas of the goal is one of the best predictors of success when executing the penalty kick [9,12]. Therefore, very often soccer players decide to kick the ball emphasizing accuracy over power. For example, during the World Cup in 2006, over 90% of penalty kicks were taken using the side-foot technique, which is a shooting technique that is used for accuracy rather than power [13].

The current study aimed to identify key factors that had an immediate influence on penalty kick accuracy. Consistent with the predictions of the OPTIMAL theory, we hypothesized that autonomy support in addition to using an external focus of attention would provide additive benefits for penalty kick accuracy compared to participants only using an external focus of attention or only being provided autonomy. Additionally, we predicted that autonomy supported participants using an external focus of attention would perform significantly better than self-controlled participants that directed their attention internally, or yoked participants that directed their attention internally. We also used a control condition to provide a baseline measure allowing us to examine the potential enhancing or depressing behavioral effects of the above experimental conditions. In addition to measuring penalty kick accuracy, a second purpose of the present study was to investigate if the self-efficacy of skilled soccer players differed as a result of the aforementioned performance conditions. It is well established that self-efficacy is critical for many team sports [14]. Therefore, we decided to measure which practice conditions produced the highest level of self-efficacy, which is defined as a person’s judgment or belief in his or her own ability to successfully execute a specific task [15]. We predicted that combining an external focus of attention with autonomy support would result in the highest amount self-efficacy compared to all other conditions.

## Material and Methods

A power analysis conducted using pilot data indicated that a minimum of 12 participants per condition were needed. To ensure sufficient power, we recruited a sample of 120 male college aged students from a larger sample of 350 students who were completing a minimum of 60 hours of soccer coaching education classes as part of a university physical education program. As part of the coaching education coursework, participants practiced the penalty kick, were required to pass a penalty kick skill test, and practiced teaching the penalty kick to others. Additionally, volunteers had to meet the following criteria: have a minimum of one year experience on an amateur soccer team, and no orthopedic injury in the past 6 months. Volunteers (mean age = 21.7 years, *s* = 1.4 years) were considered moderately skilled in the penalty kick. Women were excluded from the study. In the participant pool, 111 of volunteers were right-legged and nine were left-legged. Participants were asked to abstain from any strength and conditioning training for a minimum of three days prior to their involvement in the study. All participants signed an informed consent. The methods used in the present study were approved by a university’s Ethics Committee.

The experiment was conducted in an indoor climate-controlled sports facility. The outline of a regulation size soccer goal with four targets (i.e., black circle with a diameter of 30 cm) were placed on a white well lit wall inside the sports facility (Fig 1). A black spot representing the start location for each penalty kick was located on the floor. All penalty kicks were taken from a regulation distance of 11.00 m. Participants used a regulation soccer ball (i.e., size 5) during all penalty kicks. The locations of black circular targets inside the goal area were chosen based on previous research [12,16,17], which showed that these areas had a very high probability of successful goal scoring.

**Fig. 1.**
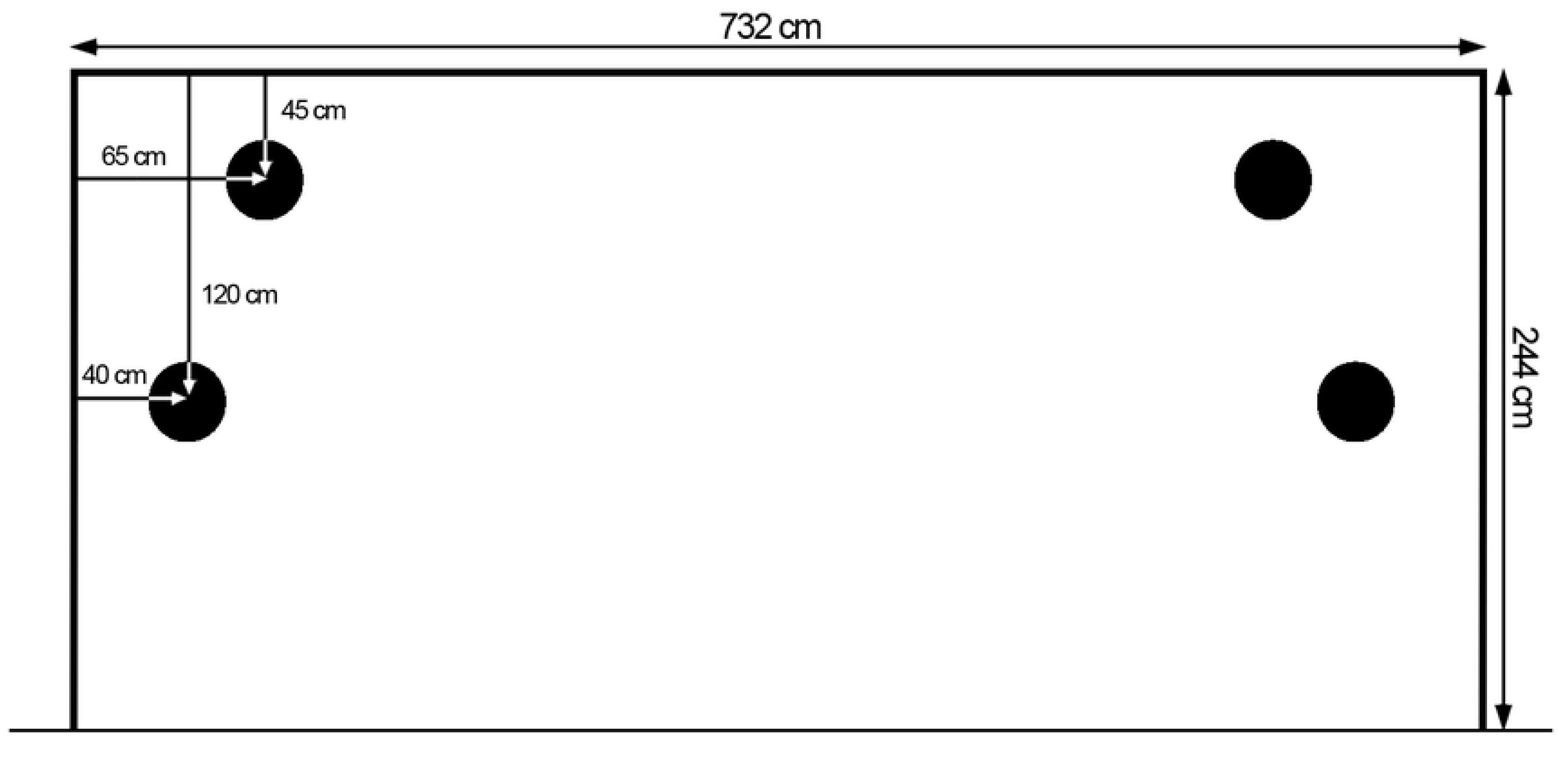
Schematic representation of the goal area and four circular targets used during the penalty kick (frontal view)

Each penalty kick was recorded by two cameras (Panasonic HC-V770). Both cameras were positioned 10 m in front (i.e., facing) of the goal. One camera was positioned 3 m to the left (camera 1) and the other was placed 3 m to right (camera 2) of the line perpendicular to the middle of the goal. The video cameras, recording at a rate of 50 Hz, were used to record the location of the ball as it hit the wall. Screen shots from the video were taken using Kinovea System® software to measure the distance between the target center and the center of the soccer ball. The height of the goalpost was used for calibration. Kicking accuracy (i.e., mean radial error) was taken as the sum of target-to-ball distances at all kicks for one participant and divided by the number of trials [18]. All penalty kicks that struck the wall within the video frame of view were included in the analysis. When a shot landed outside the camera viewing area (i.e., beyond a distance of 2.5 m from the centre of the target), that trial was scored at a distance of 2.5 m. It is important to note that only 29 shots among the total of 1440 taken shots landed outside the measurement area. In other words, 98% of the shots taken landed inside the measurement field.

In addition to performing penalty kicks, participants were asked to answer three questions regarding their self-efficacy, which was measured by a task-specific scale from 10 (not confident at all) to 100 (very confident). This instrument was adopted based on guidelines presented in Bandura’s *Guide for Constructing Self-Efficacy Scales* [19]. The same scale has been used in previous research as a reliable (Cronbach’s alpha = .96) and valid measure of self-efficacy [20]. All participants answered one question prior to performing penalty kicks (e.g., How confident are you in performing the penalty kick?) and two questions following the test (e.g., How confident were you in your ability to follow the instructions?, and How confident were you in your ability to successfully hit the correct target?). The two follow-up questions were averaged for the analysis [20].

All participants were familiarized with the aim of this study without providing any knowledge about the expected effects of attentional focus or autonomy support on penalty kick performance. Volunteers were randomly assigned to one of six groups of 20 subjects: external focus with autonomy support (EF/AS), external focus (EF), internal focus with autonomy support (IF/AS), internal focus (IF), autonomy support (AS) and control (C). Participants were instructed to perform penalty kicks with the intent of striking one of the four specified targets inside the goal area. All participants were given the following explicit instructions prior to the initiation of the experiment “hit the target as accurately as possible.” Each participant completed a total of 12 shots, with the constraint that there were three shots at each of the four targets. All shots were performed with the dominant leg and participants were allowed to use two run-up steps during each kick.

Participants in the EF/AS group were instructed to focus their attention on the given target, at the same time they were told that they could choose the order of the targets in which to kick the ball towards with the additional constraint that they could not kick the ball to the same target twice in a row. Before each shot, participants from the autonomy support groups verbally informed the experimenter about the selected target, (e.g. “I choose to kick the ball to the top target on the right side”). The EF group participants were instructed to focus their attention on the target, but the order of kicking targets for participants in this group were yoked with a counterpart in the EF/AS group. Specifically, the order of kicking targets for each person in the EF group was determined (i.e., yoked) by a paired participant in the EF/AS group. Participants in the IF/AS and IF group were asked to concentrate on the movement of the kicking leg. Participants in the IF/AS group verbally reported which target they selected prior to each kick. The order of kicking targets for the IF participants were determined by a yoked participant in the IF/AS group. Participants in the AS group were simply told to choose the order of the targets. Participants in the AS group were not provided any focus of attention instructions. Similarly, volunteers in the C group did not receive any attentional-focus instruction. The order of kicking targets for participants in C group was determined by a yoked participant in the AS group. Participants in the EF, IF and C groups were informed which target they should kick the soccer towards prior to each attempt. Experimental instructions were provided to each participant prior to the first trial and again prior to the third, sixth, ninth, and twelfth trial. Participants were not provided any feedback regarding their performance for the duration of the experiment. After each shot, the ball was recovered by the researcher and returned to the penalty kick start location. Each testing session was preceded by a warm-up consisting of a 5-minute jog and 5-minute dynamic stretching routine. Then participants took four familiarization penalty kicks aiming at each of the four circular targets. This was followed by the initiation of the main experiment.

All data are presented as means ± *SE*. Data were tested for normality and homogeneity of variance assumptions. A between-subject design was used to compare the results of the experimental and control groups. Separate one-way ANOVAs were used to determine if there were any significant differences between groups regarding kicking accuracy or self-efficacy. If main effects were observed, Tukey post-hoc tests were used for follow-up analysis. Effects sizes were estimated by two measures. First, partial eta squared (η_p_^2^) was interpreted as a small (η_p_^2^=0.01), moderate (η_p_^2^=0.06) and large effect (η_p_^2^=0.14). Second, Cohen’s d for pairwise comparisons was used based on the criteria of trivial (d=0–0.19), small (d=0.20–0.49), medium (d=0.50–0.79) and large (d=0.80 and >0.80) [21]. The alpha level was set at 0.05 for all tests. Statistica v. 13.1 software was used for all statistical calculations.

## Results

### Accuracy

Average mean radial error results for each group are presented in Figure 2. The analysis of kicking accuracy, expressed as mean radial error (Fig. 2), revealed a main effect for Group, *F*(5,114) = 5.52, *P* < 0.001, *η* _*p*_^2^ = .20. Post-hoc testing showed the EF/AS group had better kicking accuracy (i.e. closer to the target) compared to the IF/AS (*P* < 0.01, d = 1.16), IF (*P* < 0.05, d = 0.88) and C groups (*P* < 0.05, d = 0.88). Also, the EF group demonstrated better kicking accuracy compared to the IF/AS (*P* < 0.05, d = 1.37), IF (*P* < 0.05, d = 0.96) and C groups (*P* < 0.05, d = 0.98). No other group differences were observed.

**Fig. 2.**
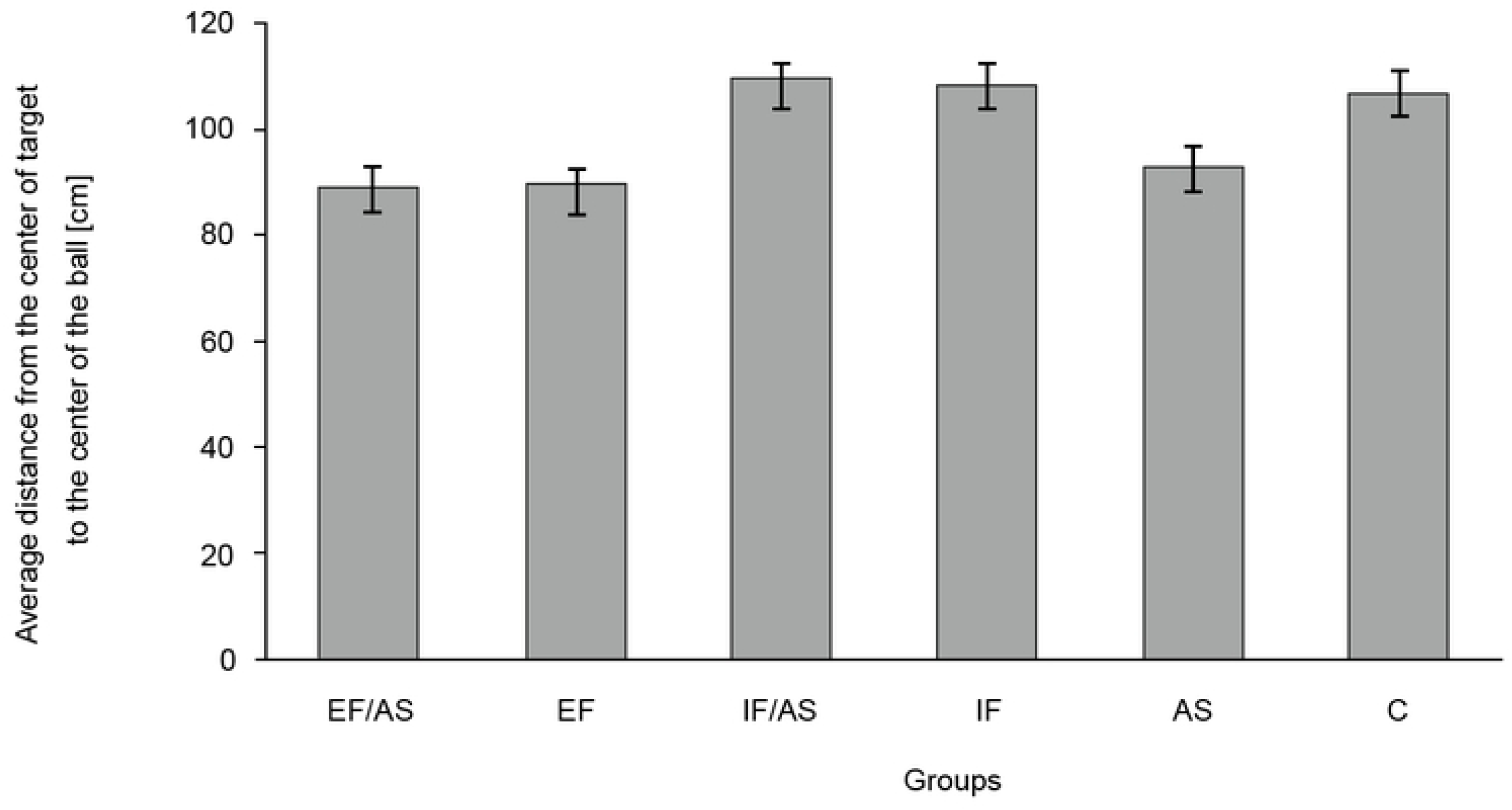
Average mean radial error for the external focus/autonomy support (EF/AS), external focus (EF), internal focus/autonomy support (IF/AS), internal focus (IF), autonomy support (AS) and control (C) groups in penalty kick performance (mean ± standard error)

### Self-efficacy

The initial measurement of self-efficacy did not reveal any significant differences between any of the experimental conditions, *F*(5,114) = 0.19, *P* > 0.05, *η*_*p*_^2^ = 0.008. However, our analysis did reveal significant differences in self-efficacy following testing, *F*(5,114) = 3.81, *P <* 0.001, *η* _*p*_^2^ = 0.14 (Fig. 3). Follow up analysis indicated that the EF/AS group demonstrated higher self-efficacy compared to the EF (*P* < 0.05, d = 1.29), IF/AS (*P* < 0.01, d = 1.24), IF (*P* < 0.05, d = 0.87) and C (*P* < 0.01, d = 1.15) groups.

**Fig. 3.**
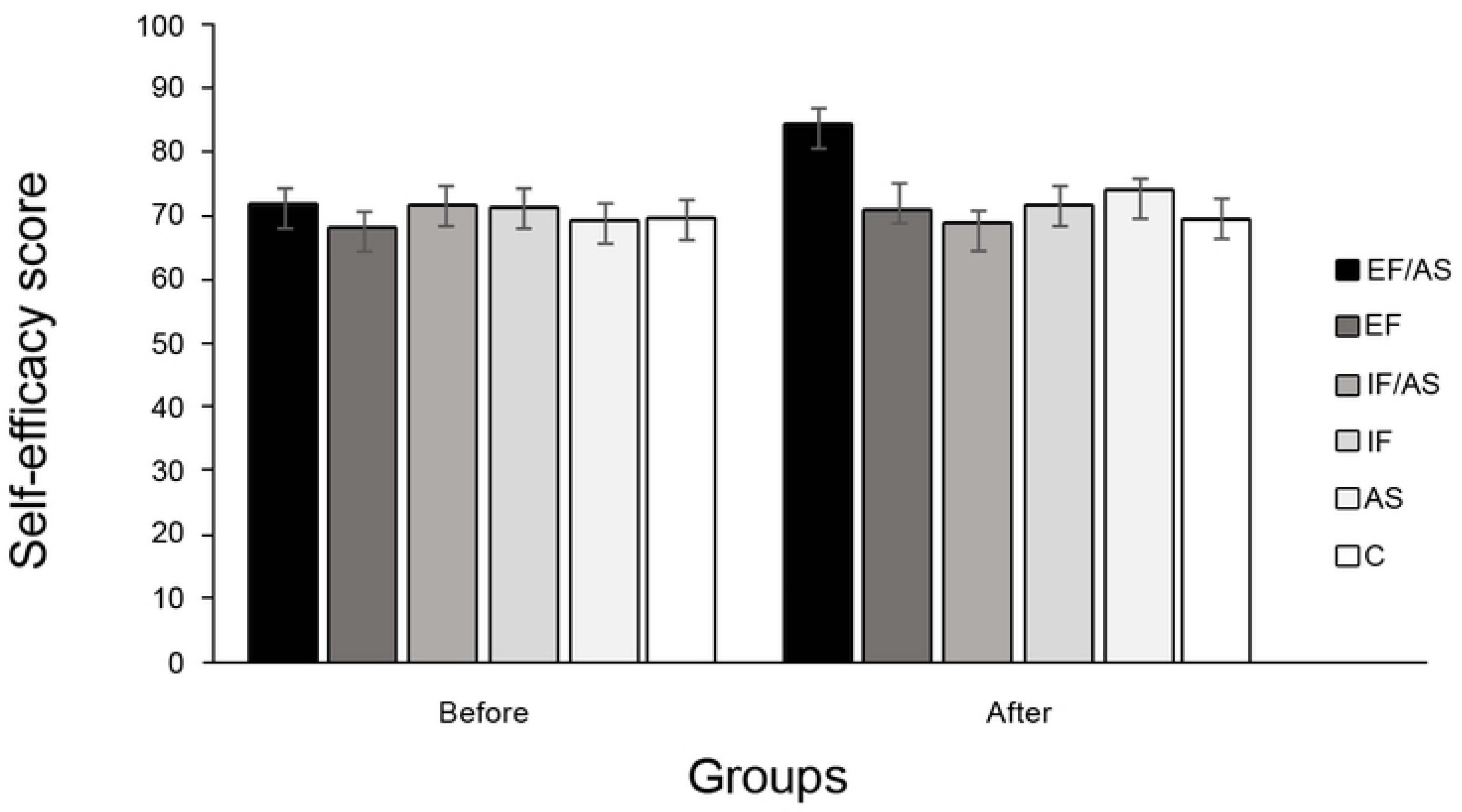
Means and standard errors of self-efficacy scores for the external focus/autonomy support (EF/AS), external focus (EF), internal focus/autonomy support (IF/AS), internal focus (IF), autonomy support (AS) and control (C) groups before and after penalty kick performance

## Discussion

The aim of this study was to investigate if the combination of an external focus of attention and autonomy facilitated a motor performance advantage compared to conditions where both factors were used independently, as predicted by the OPTIMAL theory [2]. In addition, we were interested in measuring if self-efficacy differed between the conditions as a result of practice conditions. We chose the penalty kick because it is one of the most profitable skills in sport [6].

The results of our experiment confirmed that combining an external focus of attention and autonomy support immediately improved motor performance relative to baseline trials completed in the C condition. However, our findings further revealed that these benefits were not additive as reported by Wulf et al. [5]. They suggested that combining an external focus with autonomy support showed better throwing accuracy on retention and transfer tests than did each condition alone. We did not observe such results in the present study. Here we will discuss two possible reasons why our findings differ from those reported by Wulf et al. [5]. First, in the current study we were only interested in examining the immediate effects on motor performance within a moderately skilled population performing a well-practiced task, whereas Wulf et al. [5] research examined motor learning within a low skilled population performing a novel task. When the findings of our study are compared to those reported by Wulf et al. [5], it appears there may be a skill level interaction which may impact the potential additive effects of combining an external focus of attention with autonomy support. That is, beginners may benefit by the combination of an external focus with autonomy support. In contrast, skilled performers may not display performance enhancements from this same practice strategy. The second possible reason our results differed from those reported by Wulf et al. [5] is related to the relative complexity of the tasks that were practiced in each experiment. In the Wulf et al. [5] experiment novices practiced throwing a ball with the non-dominate hand, while we tested a more complex skill (i.e., kicking) requiring more control of the action because of using more degrees of freedom relative to throwing. Therefore, the accuracy and motor control demands placed on the mover are is greater when kicking rather than throwing a ball. This is in line with Van den Tillaar and Ettema’s [22] and Van den Tillaar and Ulvik [23] research which showed that overarm throwing accuracy is easier to establish than kicking. As a result of these increased demands placed on the mover when kicking the soccer ball, it is possible that combing an external focus of attention with autonomy support did not provide additive effects. Our findings indicate that only instructing the mover to focus externally or giving them autonomy was equally effective to the combination of those two factors. Whereas beginners practicing a novel and less demanding motor skill do benefit from the combination of an external focus of attention and autonomy, as reported by Wulf and colleagues [5]. The results of our analysis indicated that the EF/AS and EF groups achieved better kicking accuracy (the ball directed closer to the target) than the control group (C), while there was no difference in accuracy between the AS and C groups. This finding indicates that the combination of external focus and autonomy support or only providing an external focus immediately improved shooting accuracy within skilled soccer players. The enhancement for the EF/AS condition provides support for the basic assumption of the OPTIMAL theory. However, the EF/AS and EF groups produced a similar level of accuracy, which does not support the predictions of the OPTIMAL theory. We propose this finding suggest that focusing attention externally has a larger impact on motor performance compared to giving the mover autonomy when performing the penalty kick. These results are not unexpected, since according to previous research examining the constrained action hypothesis [24], an external focus of attention has a strong positive influence on many skills requiring kicking accuracy. For example, the effects of attentional focus on kicking were examined by Zachry [25]. Participant in that experiment performed a place kick that is commonly used in American football. The goal of the task was to hit a target in the center of a net from a distance of 5 m. The results revealed that the external focus condition provided significantly better kicking accuracy compared to the control and internal focus conditions. Zachry [25] concluded this was due to an external focus promoting automaticity allowing the motor control system to self-organize more naturally [26], without overloading the nervous system and thus resulting in a more accurate kicking action. Previous research examining the OPTIMAL theory also showed that an external focus of attention contributes to goal-action coupling by shifting concentration towards the task goal while at the same time reducing self-focus [27]. Furthermore, it has been observed when players performed an accurate penalty kick, they fixed their attention on the desired target [28] or edge of the goal [29]. In addition to utilizing an external focus, it appears that fixating on such cues provides visual information about target location that is essential to control the action. These collective findings suggestion that in order to perform the penalty kick accurately, it is paramount that the athlete focus attention externally on the desired shot placement.

A noteworthy observation from our study is that we did not find any significant performance differences between the self-control and yoked groups. Previous studies have shown that allowing learners to control some aspect of the practice condition resulted in more effective behavioral outcomes relative to not providing them any choice [30]. There are many possible explanations for these positive results such as increasing task interest and motivation to learn, and promoting greater movement control or higher self-efficacy. However, there are only a few studies that have demonstrated beneficial effects of autonomy support on practice performance and not on motor learning [27,31]. For example, greater impact of force and velocity of punches have been demonstrated within an autonomy support condition in elite male kickboxers compared to conditions without the possibility choosing the order of punches [32]. Interestingly, some studies report that during the first trial of motor learning, self-control and yoked groups demonstrated similar performances [5,33]. Using these observations, we conjecture that practice performance may be less sensitive for beneficial outcomes of autonomy support conditions than of those in motor learning.

The findings of previous research have indicated [2] that providing a mover with autonomy influences motivation, including self-efficacy. In the current study, participants across all conditions reported a similar level of self-efficacy prior to the testing session. However, after the completion of the 12 penalty kicks we found that the EF/AS group reported a higher level of self-efficacy compared to the EF group. This is in line with findings reported by Wulf et al. [5], where individuals with task-relevant autonomy support resulted in higher self-efficacy compared with conditions which did not offer choices. These findings are also in agreement with the OPTIMAL theory. Specifically, our findings demonstrated that providing an external focus of attention with autonomy effected self-efficacy within our moderately skilled sample. Another factor that is important to mention here is that successfully performing a penalty kick is a very stressful task [34]. Previous researchers [14] have proposed that effective practice should foster mechanisms that protect the mover against stress. Therefore, we believe that having a high level of self-efficacy as a result of directing attention externally with autonomy support positively influenced the performance of the stressful task of successfully executing the penalty kick. This possibility needs to be addressed in future research by including a goalkeeper that is attempting to block the shot. The addition of a goalkeeper would increase the stressful nature of the task. If the combination of an external focus of attention with autonomy support does in fact protect the mover from stress, then such a form of practice should result in more accurate kicks relative to movers that practiced with only an external focus of attention or with autonomy.

## Conclusion

The current study showed that an external focus of attention and autonomy support are associated with overall better motor performance. Combining an external focus of attention with autonomy support immediately enhanced penalty kick accuracy relative to participants that performed in the control condition and those that received a combination of an internal focus of attention with autonomy support and their yoked counterparts. Although we did not find additive advantages for motor performance comparing the external focus and autonomy support to external condition without a motivational manipulation. However, we did see that combining an external focus of attention with autonomy support did significantly improve self-efficacy compared to all other conditions. In summary, we confirmed that matching attentional and motivational factors optimized the performance of the penalty kick. The present results also suggest that self-control practice involving the selection of the desired target led to better accuracy during penalty kick performance relative to instructions which directed attention internally or a form of practice that combined an internal focus with autonomy support. In agreement with earlier findings and the OPTIMAL theory, the findings reported here support that attentional and motivational variables directly influence motor performance and self-efficacy in moderately skilled soccer players.

## References

1. Wulf G. Attentional focus and motor learning: a review of 15 years. Int Rev Sport Exerc Psychol. 2013; 6(1):77–104. https://doi.org/10.1080/1750984X.2012.723728

2. Wulf G, Lewthwaite R. Optimizing performance through intrinsic motivation and attention for learning: The OPTIMAL theory of motor learning. Psychon Bull Rev. 2016; 23(5):1382–414. https://doi.org/10.3758/s13423-015-0999-9. PMID: 26833314

3. Pascua LA, Wulf G, Lewthwaite R. Additive benefits of external focus and enhanced performance expectancy for motor learning. J Sports Sci. 2015; 33(1):58–66. https://doi.org/10.1080/02640414.2014.922693. PMID: 24875153

4. Wulf G, Chiviacowsky S, Cardozo PL. Additive benefits of autonomy support and enhanced expectancies for motor learning. Hum Mov Sci. 2014; 37:12–20. https://doi.org/10.1016/j.humov.2014.06.004. PMID: 25046402

5. Wulf G, Chiviacowsky S, Drews R. External focus and autonomy support: Two important factors in motor learning have additive benefits. Hum Mov Sci. 2015; 40:176–84. https://doi.org/10.1016/j.humov.2014.11.015. PMID: 25589021

6. Dalton K, Guillon M, Naroo SA. An analysis of penalty kicks in elite football post 1997. Int J Spor Sci Coach. 2015; 10(5):815–27. https://doi.org/10.1260%2F1747-9541.10.5.815

7. Franks IM, Hanvey T. Cues for goalkeepers: high-tech methods used to measure penalty shot response. Soccer J. 1997; 42:30–3.

8. McGarry T, Franks IM. On winning the penalty shoot-out in soccer. J Sports Sci. 2000; 18(6):401–9. https://doi.org/10.1080/02640410050074331. PMID: 10902675

9. Bar-Eli M, Azar OH. Penalty kicks in soccer: an empirical analysis of shooting strategies and goalkeepers’ preferences. Soccer Soc. 2009; 10(2): 183–91. https://doi.org/10.1080/14660970802601654

10. Binsch O, Oudejans RR, Bakker FC, Savelsbergh GJ. Ironic effects and final target fixation in a penalty shooting task. Hum Mov Sci. 2010; 29(2):277–88. https://doi.org/10.1016/j.humov.2009.12.002. PMID: 20206393

11. Navarro M, van der Kamp J, Ranvaud R, Savelsbergh GJ. The mere presence of a goalkeeper affects the accuracy of penalty kicks. J Sports Sci. 2013; 31(9):921–9. https://doi.org/10.1080/02640414.2012.762602. PMID: 23360203

12. Kerwin DG, Bray K. Measuring and modelling the goalkeeper’s diving envelope in a penalty kick. In: Moritz EF, Haake S, editors. The Engineering of sport 6. New York: Springer; 2006. p. 321–6.

13. Lees A, Owens L. Early visual cues associated with a directional place kick in soccer. Sport Biomech. 2011; 10(02):125–34. https://doi.org/10.1080/14763141.2011.569565. PMID: 21834396

14. Feltz DL, Short SE, Sullivan PJ. Self-efficacy in sport. Champaign IL: Human Kinetics; 2008.

15. Bandura A. Self-efficacy: toward a unifying theory of behavioral change. Psycho Rev. 1977; 84(2):191. https://doi.org/10.1037/0033-295X.84.2.191

16. Graham-Smith P, Lees A, Richardson D. Analysis of technique of goalkeepers during the penalty kick. J Sports Sci. 1999; 17(11):905–29. https://doi.org/10.1080/026404199365461

17. Almeida CH, Volossovitch A, Duarte R. Penalty kick outcomes in UEFA club competitions (2010-2015): The roles of situational, individual and performance factors. Int J Perform Anal Sport. 2016; 16(2):508–22. https://doi.org/10.1080/24748668.2016.11868905

18. Hancock GR, Butler MS, Fischman MG. On the problem of two-dimensional error scores: Measures and analyses of accuracy, bias, and consistency. J Mot Behav. 1995; 27(3):241–50. https://doi.org/10.1080/00222895.1995.9941714. PMID: 12529235

19. Bandura A. Guide for constructing self-efficacy scales. In: Pajares F, Urdan T, editors. Self-efficacy beliefs of adolescents. Adolescence and education. Greenwich: Information Age Publishing; 2006; 4, p. 307–37.

20. Saemi E, Porter JM, Ghotbi-Varzaneh A, Zarghami M, Maleki F. Knowledge of results after relatively good trials enhances self-efficacy and motor learning. Psychol Sport Exerc. 2012; 13:378–82. https://doi.org/10.1016/j.psychsport.2011.12.008

21. Larson-Hall J. How to run statistical analyses. In: Mackey A, Gass SM, editors. Research methods in second language acquisition: A practical guide. London: Blackwell Publishing; 2012; p. 245–8.

22. Van den Tillaar R, Ettema G. Influence of instruction on velocity and accuracy of overarm throwing. Percept Mot Skills. 2003; 96(2):423–34. https://doi.org/10.2466/pms.2003.96.2.423. PMID: 12776824

23. Van den Tillaar R, Ulvik A. Influence of instruction on velocity and accuracy in soccer kicking of experienced soccer players. J Mot Behav. 2014; 46(5):287–91. https://doi.org/10.1080/00222895.2014.898609. PMID: 24773185

24. Wulf G, McNevin N, Shea CH. The automaticity of complex motor skill learning as a function of attentional focus. Q J Exp Psychol. 2001; 54A:1143–54. https://doi.org/10.1080/713756012. PMID: 11765737

25. Zachry TL. Effects of attentional focus on kinematics and muscle activation patterns as a function of expertise. [master’s thesis]. Las Vegas (NV): University of Nevada; 2005.

26. Bernstein NA. On dexterity and its development. In: Latash ML, Turvey MT, editors. Dexterity and its development. Mahwah, NJ: Lawrence Erlbaum; 1996. p. 171–204.

27. Abdollahipour R, Nieto MP, Psotta R, Wulf G. External focus of attention and autonomy support have additive benefits for motor performance in children. Psychol Sport Exerc. 2017; 32:17–24. https://doi.org/10.1016/j.psychsport.2017.05.004

28. Wood G, Wilson MR. Quiet-eye training for soccer penalty kicks. Cogn Process. 2011; 12(3):257–66. https://doi.org/10.1007/s10339-011-0393-0. PMID: 21318734

29. Timmis MA, Turner K, Van Paridon KN. Visual search strategies of soccer players executing a power vs. placement penalty kick. PLoS ONE. 2014; 9(12):115–79. https://doi.org/10.1371/journal.pone.0115179. PMID: 25517405

30. Chiviacowsky S. Self-controlled practice: Autonomy protects perceptions of competence and enhances motor learning. Psychol Sport Exerc. 2014; 15(5):505–10. https://doi.org/10.1016/j.psychsport.2014.05.003

31. Englert C, Bertrams A. Autonomy as a protective factor against the detrimental effects of ego depletion on tennis serve accuracy under pressure. Int J Spor Exerc Psychol. 2015; 13(2):121–31. https://doi.org/10.1080/1612197X.2014.932828

32. Halperin I, Chapman DW, Martin DT, Lewthwaite R, Wulf G. Choices enhance punching performance of competitive kickboxers. Psychol Res. 2016; 81(5):1051–8. https://doi.org/10.1007/s00426-016-0790-1. PMID: 27465395

33. Iwatsuki T, Abdollahipour R, Psotta R, Lewthwaite R, Wulf G. Autonomy facilitates repeated maximum force productions. Hum Mov Sci. 2017; 55:264–8. https://doi.org/10.1016/j.humov.2017.08.016. PMID: 28865313

34. Bar-Eli M, Friedman Z. Psychological stress in soccer: The case of penalty kicks. Soccer J. 1988; 33(6):49–52.

